# Interdisciplinarity through internationality: results from a US-Mexico graduate course bridging computational and plant science

**DOI:** 10.1101/2024.06.19.599776

**Authors:** Daniel H. Chitwood, Alejandra Rougon-Cardoso, Robert VanBuren

## Abstract

Interdisciplinarity is used to integrate and synthesize new research directions between scientific domains, but it is not the only means by which to generate novelty by bringing diverse perspectives together. Internationality draws upon cultural and linguistic diversity that can potentially impact interdisciplinarity as well. We created an interdisciplinary class originally intended to bridge computational and plant science that eventually became international in scope, including students from the US and Mexico. We administered a survey over four years designed to evaluate student expertise. The first year of the survey included only US students and demonstrated that biology and computational student groups have distinct expertise but can learn the skills of the other group over the course of a semester. Modeling of survey responses shows that biological and computational science expertise is equally distributed between US and Mexico student groups, but that nonetheless these groups can be predicted based on survey responses due to sub-specialization within each domain. Unlike interdisciplinarity, differences arising from internationality are mostly static and do not change with educational intervention and include unique skills such as working across languages. We end by discussing a distinct form of interdisciplinarity that arises through internationality and the implications of globalizing research and education efforts.

## Introduction

A number of grand challenges, revolving around the related issues of agriculture, environmental integrity, feeding the world, and climate change (Robertson and Swinton, 2005; Hertel, 2015; Dhankher and Foyer, 2018; Ryan et al., 2018) will require coordinated contributions across science, technology, engineering, and mathematics (STEM) disciplines. Training the next generation of scientists not only requires an infusion of computational (Ruinstein and Chor, 2014) and mathematical (Maass et al., 2019) skills into fields that are foundational to address these challenges, but also to synthesize new interdisciplinary and transdisciplinary research domains and provide educational curricula for them, critically defining and assessing the aims of these novel undertakings (Gao et al., 2020). Because these grand challenges are global in scope, the research and training we undertake to address them must be equally inclusive. That scientific and mathematical talent are equally distributed across the world and have manifested in diverse ways across cultures is taken as axiomatic (Ardila-Mantilla, 2016), but global disparities in scientific participation are deeply rooted in history and persist to this day (Jeffries-EL, 2022). We highlight North American agriculture as one example. Even though the manual labor that sustains US agriculture is 54% of Mexican origin, 31% white (not Hispanic), and 44% are not US citizens (USDA Economic Research Service, 2023), the plant sciences that represent this societal sphere are overwhelmingly white and male (NCSES, 2023). Similar disparity, in those who are allowed to participate in STEM research and education versus the communities denied access who are most vulnerable to the very problems science seeks to address, is global in scale (Marks et al. 2021; 2023). While interdisciplinarity is important to address the grand challenges of our time, internationality and the lack of global connectedness in STEM fields remains a largely unaddressed concern.

Just as we can center interdisciplinary learning objectives, so too can we teach intercultural competencies (Wickenhauser and Karcher, 2020). Although the pandemic presented higher education challenges in online and remote learning (Ali et al., 2020), these platforms also invited instructors to incorporate intercultural opportunities (Wickenhauser, 2021). Massive Open Online Courses (MOOC) by their nature transcend borders and can potentially include global learning objectives by which students “[learn] about the world, [learn] with others, and [learn] to act” (Mathews and Landorf, 2016). Teaching such global competency requires self-awareness of the place and context of students, instructors, and institutions with respect to local and international communities and culturally aware practices (MacCleoud, 2018). Just as “parachute science”, in which researchers from wealthy countries extract information and resources from poorer communities without including them (Stefanoudis et al., 2021), acts against a globally inclusive research community, the balance of power within a global classroom needs to be evaluated through a critical lens. Online education needs to be mindful of cognitive, social, and teacher presence (Garrison, 2007) and so too do these processes influence interactions in an international and intercultural classroom. Despite the challenges facing teaching and learning in a global context, the benefits are potentially far-reaching, not only addressing core issues holding us back from truly addressing grand challenges, but potentially internationalizing higher education through deliberate efforts (Kahn and Agnew, 2017).

Here, we report on a class that began at Michigan State University (MSU) in the United States as an interdisciplinary effort to integrate computational and plant science learning objectives, but when shifted online during the pandemic became international in scope, offered to Universidad Nacional Autónoma de México (UNAM) students in Mexico as well. Notably, professors of both institutions in both countries participated. A survey in which students self-assess their expertise in plant and computational learning objectives given throughout the semester demonstrates unique competencies between biology and computational science student groups, and that these competencies are equally distributed among US and Mexican students. Yet, MSU and UNAM groups can be predicted from survey responses due to sub-specialization within biology and computational domains. We argue that internationality promotes diversity and interdisciplinarity in STEM through complementary expertise that arises between cultures due to unique motivations and education experiences.

## Materials and Methods

This study was determined to be exempt under the Revised Common Rule under 45 CFR 46.104(d) 1, 4ii by the Michigan State University (MSU) Institutional Review Board (IRB) on 3 April 2024 (MSU Study ID: STUDY00010594).

### Curriculum and classroom environment

The graduate class for which data was collected originated from a National Science Foundation (NSF) Research Traineeship (NRT) grant at MSU called IMPACTS (Integrated training model in Plant And Compu-Tational Sciences; #1828149; NRT-IMPACTS, 2024). There were five main training objectives to the grant: 1) Proficiency in core knowledge in plant & computational science, 2) Expertise in interdisciplinary research in plant biology & computation, 3) Development of communication, leadership, & management skills, 4) Development of trainees’ teaching skills, and 5) Development of trainees’ mentoring skills. To address the first point, a series of classes were created to teach and integrate plant biology and computational science learning objectives. The course we describe in this manuscript was the first in the series and titled *HRT841: Foundations in Computational and Plant Sciences*. The course started in 2019 and in 2019 and 2020 consisted only of MSU students. Using Jupyter notebooks (Kluyver et al., 2016), an interactive coding environment with markdown text and in which media are embedded that is used for research, scientific reproducibility, and educational purposes (and with which the analysis for this manuscript are carried out), the class introduces biology students with no prior coding experience and computational students with no prior experimental experience to introductory coding lessons, inspired by examples from plant biology, in Python. The class also teaches command line and bioinformatics. In the second half of the course, students collaborate and author a class project that applies computational approaches to a plant biology question. For example class projects, see published projects from 2019 (Bryson et al., 2020), 2020 (Palande et al., 2023a), and 2021 (Palande et al., 2023b).

When the COVID-19 pandemic began in 2020 forcing higher educational institutions worldwide to shut down, the course focus on computational approaches, and the electronic format of Jupyter notebooks with embedded YouTube video lectures that could easily be disseminated, provided an opportunity to expand our student audience. The shift to a virtual environment precipitated by the pandemic also permitted us to experiment with international education. In 2021, we expanded the course to include students from the UNAM system. The students took the course for credit in the UNAM Biological Sciences postgraduate program as *Bioinformática y minería de datos con python* and in the UNAM ENES León Agrigenomic Sciences undergraduate program as *Plantas y Python*. We published the materials online as a Jupyter book titled *Plants&Python* in English and Spanish for anyone with an internet connection to access the course (https://plantsandpython.github.io/PlantsAndPython; VanBuren et al., 2022; Williams, 2022). Using a flipped classroom approach, students used the online educational materials and worked together in groups during class on a Discord server. After the pandemic, we have continued to use a hybrid approach, with MSU students optionally attending class in person with instructors, Mexican students participating virtually, and the online Discord server serving as a communication channel and digital archive for the yearly class project.

### Survey

In 2020 (the second year of the course) we began administering a survey to evaluate the degree that dueling plant biology and computational science learning objectives were being effectively taught. The survey was given before, at the middle, and end of the semester and asked students to self-assess their expertise across biology and computational science topics. In 2020 there were only MSU students that were enrolled in plant biology or computational/data science programs that were easily classified into “biology” and “computational” groups. For the remaining years of the survey (2021 to 2023), we also included students from the UNAM system. Students from throughout Mexico were enrolled in diverse academic programs (for example, animal science, chemistry, epidemiology, medical and veterinary sciences, evolutionary biology, etc.) and could not be classified into “biology” and “computational” groups easily. For the years 2021 to 2023, students were assigned to “MSU” and “UNAM” groups. We added a few more questions to the survey beginning in 2021 that asked about new learning objectives as well as working across languages and countries.

Survey questions asked students to self-assess their expertise on a 1 to 10 scale for the following categories: 1) coding, 2) statistics, 3) modeling, 4) bioinformatics, 5) computational resources, 6) molecular biology, 7) genetics & breeding, 8) plant development, 9) phylogenetics, 10) working on a team, 11) project management, 12) interdisciplinary science, 13) science communication, and 14) scientific writing. Three additional questions were added for the years 2021 to 2023: 15) command line, 16) working across countries, and 17) working across languages.

### Modeling and data analysis

Using Linear Discriminant Analysis (LDA) we created two different classification models. LDA models a categorical variable as a function of numerous continuous variables. For the number of modeled categorical variable groups *n*, LDA returns *n-1* discriminant axes that maximize the differences between groups. In this study we always model two groups, which returns one discriminant axis, which conveniently summarizes modeled group differences based on numerous survey questions as a single value. The weighted contributions of the variables to the classification are known as scalings and provide insights into how the model is working and the most important contributing factors that comprise it. Once an LDA model is created, new data can be projected onto the model and classified. The first model classifies students as belonging to biology or computational groups and is built using only 2020 data from MSU students. We then use this model to classify MSU and UNAM students as belonging to biology or computational groups for the years 2021 to 2023. The second model uses only data from 2021 to 2023 and classifies students as belonging to MSU or UNAM groups.

Data was imported using the pandas module (McKinney, 2010) as a dataframe. Masking was used to isolate specific factor levels as needed. Data scaling, LDA modeling, and cross-validation of models were performed using the scikit-learn (Pedregosa et al., 2011) functions StandardScaler, LinearDiscriminantAnalysis, and RepeatedStratifiedKFold functions, respectively. Significant differences between Linear Discriminant scores were determined using a Kruskal-Wallis test with the stats.kruskal function from the scipy (Virtanen et al., 2020) stats module. All analyses and visualizations were performed in Python using numpy (Harris et al., 2020), matplotlib (Hunter, 2007), and seaborn (Waskom, 2021) modules.

## Results

We first modeled survey responses of self-assessed student expertise as a function of whether the student was enrolled in a biology or computational program (**Figure 1A**). We built the model using only 2020 data from MSU students which could easily be classified as “biology” or “computational”. Because the course was not yet international at this time, our main objective was to evaluate the initial self-assessed expertise of students in an interdisciplinary course and monitor how this shifted during the semester. Linear Discriminant Analysis (LDA) revealed strong differences in student responses between the two groups overall (p value = 8.24 x 10^-7^, **Table 1**). The average cross-validated accuracy of the model was 93.6%. Biology students (associated with negative linear discriminant values) were more likely to express expertise in biology-related topics, as reflected in the scaling values of the survey questions (the questions with the most negative scaling values were “molecular biology”, “plant development”, and “coding”; **Figure 1B**). Computational students (associated with positive linear discriminant values) were more likely to express expertise in quantitative- and computational-related topics (for example, “statistics”, “modeling”, and “phylogenetics”). As the semester progressed, biology student responses became more computational-like (p value = 0.022) and likewise computational student responses became more biology-like (p value = 0.037; **Table 1**). We interpreted this result positively with respect to the original interdisciplinary goal of the course: that biology and computational students would mutually increase their familiarity with and expertise in learning objectives of the other group.

**Table 1:** p values for linear discriminant value comparisons for the biology vs. computational student model.

| year | comparison | p value |
| --- | --- | --- |
| 2020 | bio. / comp. | 8.24 x 10<sup>-7</sup> |
| 2020 | within bio. | 0.022 |
| 2020 | within comp. | 0.037 |
| 2021 | MSU / UNAM | 0.81 |
| 2021 | within MSU | 0.29 |
| 2021 | within UNAM | 0.41 |
| 2022 | MSU / UNAM | 0.46 |
| 2022 | within MSU | 0.72 |
| 2022 | within UNAM | 0.99 |
| 2023 | MSU / UNAM | 0.58 |
| 2023 | within MSU | 0.78 |
| 2023 | within UNAM | 0.15 |

**Figure 1:**
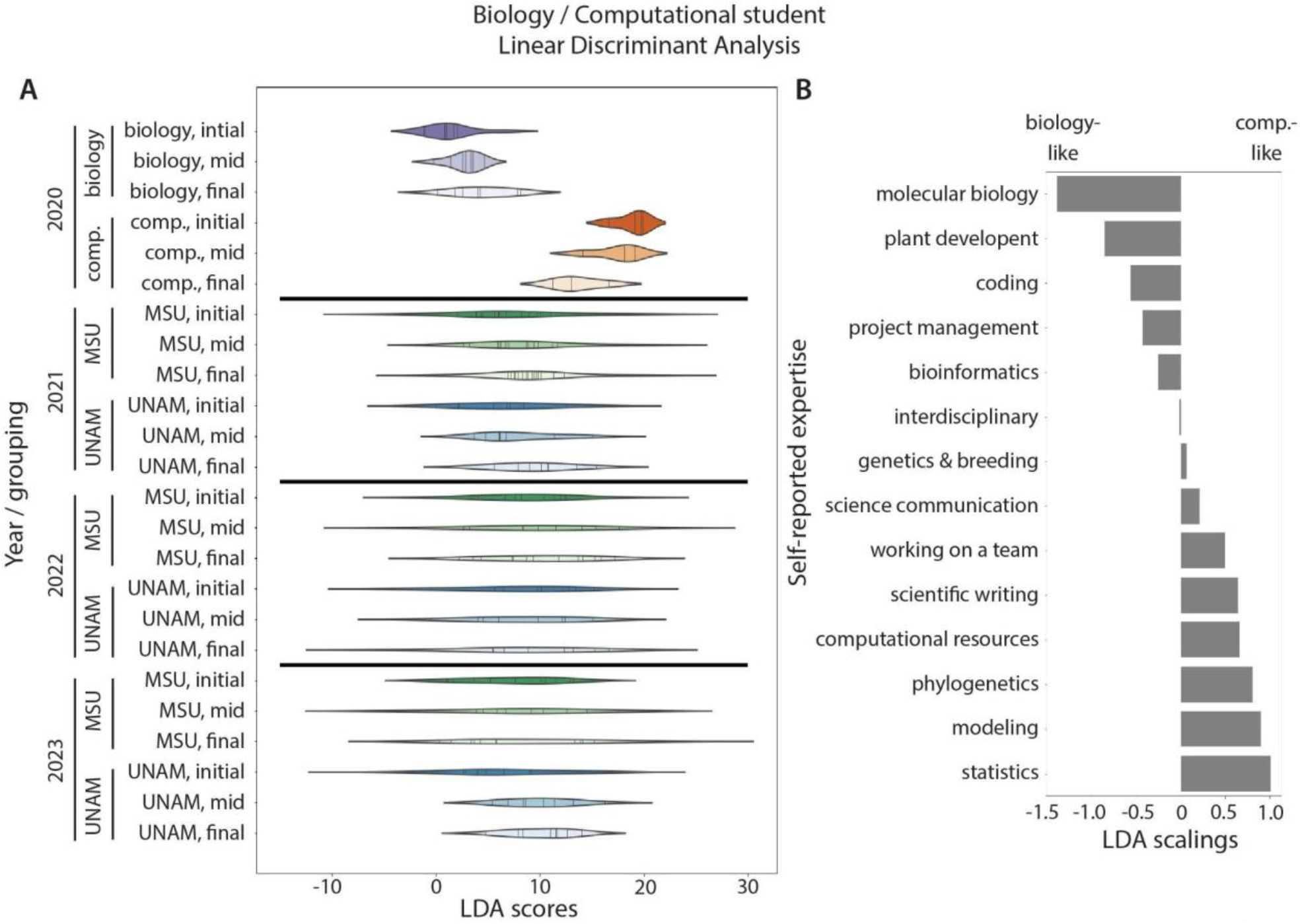
Biology vs. computational student LDA. **A)** An LDA model was constructed for biology vs. computational student identity as a function of MSU student survey responses in 2020. Linear discriminant scores for initial, mid, and final semester survey responses were calculated from the model for biology and computational student groups in 2020 and MSU and UNAM student groups 2021 to 2023. **B)** Scalings for linear discriminant scores. More negative scaling values indicate a survey question that contributes towards negative linear discriminant scores, and likewise more positive scaling values indicate survey questions that contribute to positive linear discriminant scores.

In 2021 to 2023, we included UNAM students from throughout Mexico in the class in addition to MSU students. We continued to administer the same survey with a few additional questions. Because of the diversity of academic programs offered by UNAM, we could not classify students as “biology” or “computational”. However, using the 2020 LDA model, we can predict whether student survey responses are more biology- or computational-like. Projecting MSU and UNAM student responses for 2021 to 2023 onto the 2020 model, linear discriminant values are highly variable and centered between “biology” and “computational” student groups from 2020 (**Figure 1A**). There is no statistical difference in discriminant values between MSU and UNAM groups for all years, and within each group for each year, there is no statistical change in survey responses during the semester (**Table 1**). We conclude that predicted biology and computational self-assessed expertise is equally distributed between MSU and UNAM student groups.

Although “biology” and “computational” expertise were equally distributed across MSU and UNAM groups, we wanted to determine if their survey responses from each other differed. We therefore modeled survey responses as a function of whether the student was from MSU or the UNAM system (**Figure 2A**). We built the model using all data from 2021 to 2023 (excluding 2020 data from only MSU students). In 2021, 2022, and 2023 there were strong differences between MSU and UNAM students overall (p value = 3.58×10^*-9*^, 3.77×10^-9^, 2.39×10^-8^, respectively; **Table 2**). The average cross-validated accuracy of the model was 78.0%. Besides in 2021 when MSU student responses became more UNAM-like as the semester progressed (p value = 0.012), there were no significantly different changes in responses within MSU or UNAM groups for any year (**Table 2**). Although survey questions were originally designed to assess student expertise in biology and computational disciplines, and such expertise is equally distributed between MSU and UNAM groups (**Figure 1**; **Table 1**), a combination of survey responses nonetheless is able to predict MSU and UNAM student identities. Looking at scaling values (**Figure 2B**), the negative linear discriminant values that predict MSU student identity are associated with survey questions such as “genetics & breeding”, “coding”, and “project management”, whereas positive linear discriminant values that predict UNAM student identity are associated with “molecular biology”, “working across languages”, and “bioinformatics”. We conclude that although biology and computational expertise is equally distributed across MSU and UNAM student groups, that each group specializes in sub-disciplines within each category, and that specific expertise can be associated with a single group (for example, “working across languages” with UNAM students).

**Table 2:**
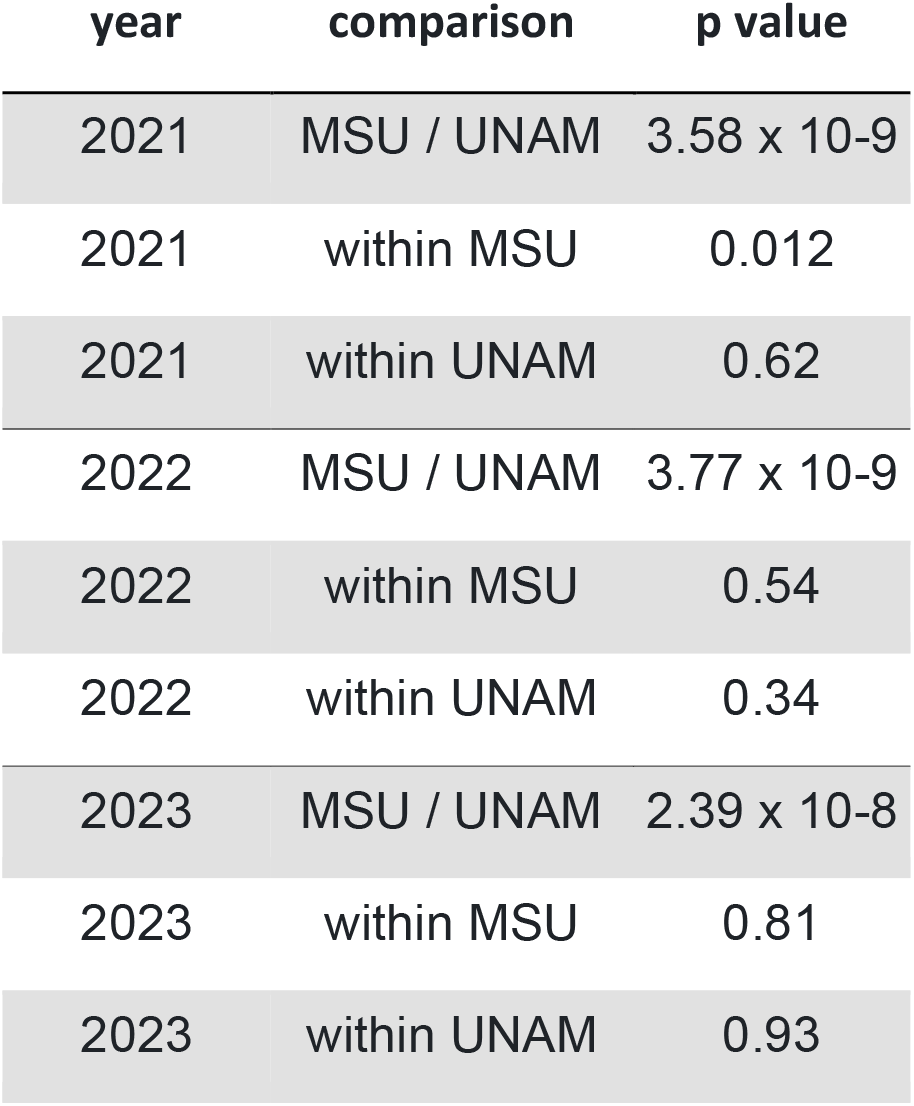
p values for linear discriminant value comparisons for the MSU vs. UNAM model.

| year | comparison | p value |
| --- | --- | --- |
| 2021 | MSU / UNAM | 3.58 x 10<sup>-9</sup> |
| 2021 | within MSU | 0.012 |
| 2021 | within UNAM | 0.62 |
| 2022 | MSU / UNAM | 3.77 x 10<sup>-9</sup> |
| 2022 | within MSU | 0.54 |
| 2022 | within UNAM | 0.34 |
| 2023 | MSU / UNAM | 2.39 x 10<sup>-8</sup> |
| 2023 | within MSU | 0.81 |
| 2023 | within UNAM | 0.93 |

**Figure 2:**
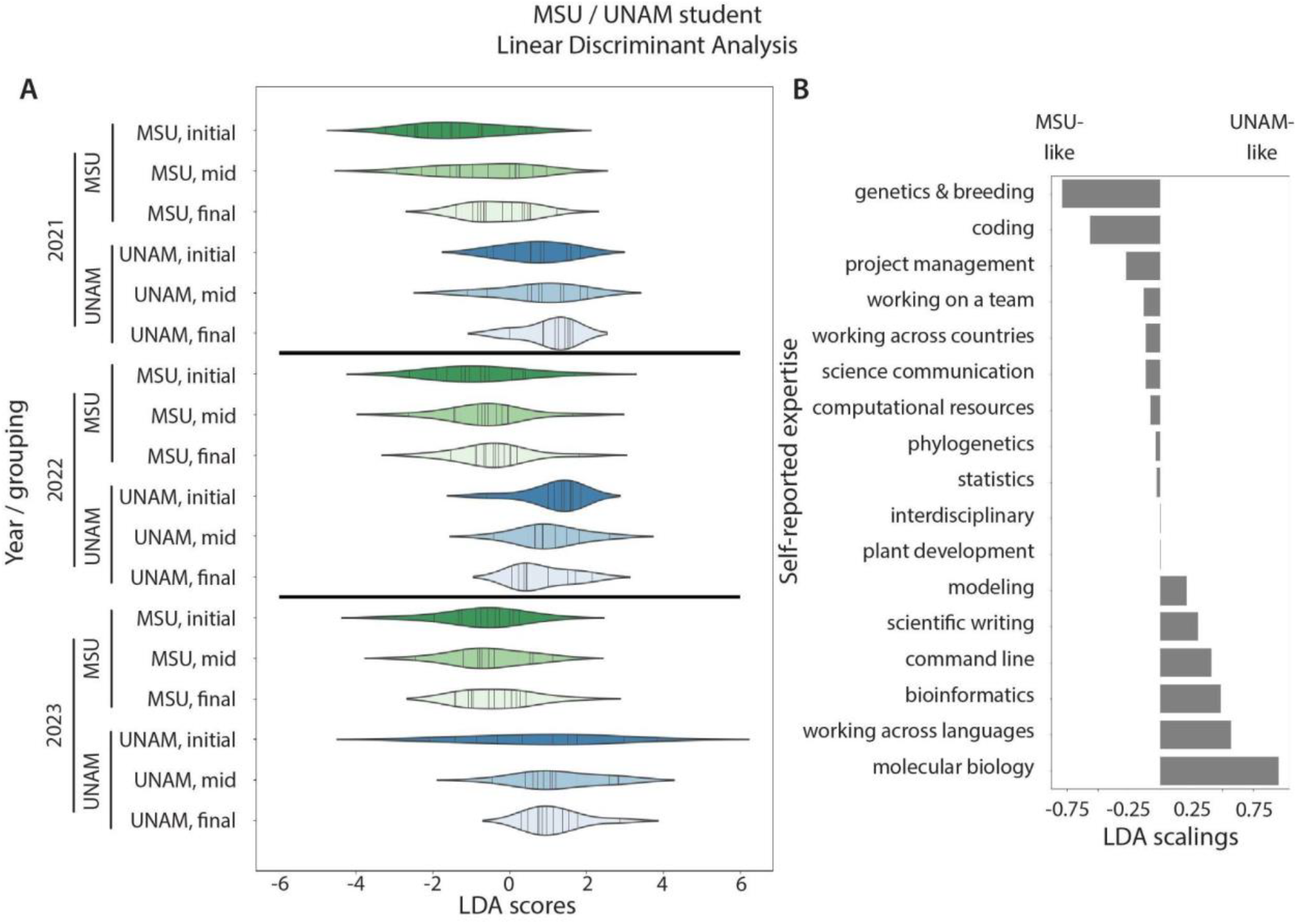
MSU vs. UNAM student LDA. **A)** An LDA model was constructed for MSU vs. UNAM student identity as a function of student survey responses from 2021 to 2023. Linear discriminant scores for initial, mid, and final semester survey responses were calculated from the model for MSU and UNAM student groups 2021 to 2023. **B)** Scalings for linear discriminant scores. More negative scaling values indicate a survey question that contributes towards negative linear discriminant scores, and likewise more positive scaling values indicate survey questions that contribute to positive discriminant scores.

## Discussion

Our course was originally designed to be interdisciplinary, bringing together plant and computational science disciplines. The course was designed at MSU, and the academic structure of the university influenced the curriculum, instructors, participants, and outcomes. The survey results we analyze in this study were designed to evaluate the efficacy of teaching plant and computational science learning objectives to students from each of these groups, but the epistemological impact of this endeavor arising exclusively from MSU cannot be overstated. In this regard, the original structure and intended impacts of our evaluative research are similar to other interdisciplinary programs.

At its heart, interdisciplinarity is about bringing two different disciplinary groups together, each with different topical interests, but also distinct cultures, perspectives, and motivations. Both the pandemic and inclusion of UNAM students into the course were unexpected and as we have already stated and the diversity of disciplines represented among UNAM students from throughout Mexico defied categorization into “biology” and “computational” groups. The influence of including students from academic institutions in a different country was so great that the originally intended structure of our interdisciplinary course across scientific domains was replaced with international designations of “MSU” and “UNAM”. If interdisciplinary research derives its benefits from the synergy of disparate domains of knowledge, then internationality potentially does the same, mixing groups with not only potentially differing academic specialization, but worldviews arising from social, cultural, and linguistic diversity.

Both interdisciplinarity and internationality bring two groups together, but our results suggest that they are not equivalent. Interdisciplinarity between “biology” and “computational” groups is defined by strong differences in self-reported expertise (**Figure 1**). Importantly, these differences can be mitigated by training. Biology students became more computational-like in their survey responses and vice versa as the semester progressed (**Table 1**). Internationality is distinct. To begin, internationality is orthogonal to interdisciplinarity, in that predicted “biology” and “computational” student groups are equally distributed among MSU and UNAM students (**Figure 1**). The differences between MSU and UNAM groups are much weaker than those between “biology” and “computational” groups (**Figure 2**), but except for 2021 MSU students which became more UNAM-like, are static and unchanging (**Table 2**). This weaker distinction between US and Mexican students, embedded within an equal distribution of biological and computational science expertise, could reflect fundamental differences in training between the countries, epistemological differences in knowledge production, differing motivations, or otherwise intrinsically different ways that MSU and UNAM students interact within broader domains. This is not unexpected, considering the profound effects of language and culture on how we interact with the world that transcend disciplinary differences. It is worth noting that while most of the differences arise through complementary, sub-specialization (for example, MSU students report more expertise in genetics and breeding and coding while UNAM students more expertise in molecular biology and bioinformatics; **Figure 2B**), some of these differences are asymmetric and unique. Notably UNAM students, who largely as native Spanish speakers were participating in a mostly English class as a second language, uniquely reported “working across languages” as an expertise.

## Conclusions

Although the scientific community is strongly focused on interdisciplinary approaches to bridge differences between domains to address global grand challenges, it is not the only way to diversify research. We show that internationality adds a distinct facet of interdisciplinary expertise, distinct from interdisciplinarity itself, and that international students provide unique skills, such as working across languages, that are otherwise missing in a local setting. Our results suggest that international education can enhance inter- and trans-disciplinary research and is an underutilized means of intentionally globalizing research to address grand challenges in an era when virtual technologies enable it.

## Data availability

The code and data necessary to reproduce the results in this manuscript are available in the following GitHub repository: https://github.com/DanChitwood/interdisciplinary_international_education

## Acknowledgements

We thank biology and computational students from Michigan State University and the Universidad Nacional Autónoma de México system for taking the *Plants&Python* course, engaging with the class, and taking time to complete the survey. We thank Tammy Long and Jyothi Kumar for organizing valuable evaluative feedback about the course and Addie Thompson and Shinhan Shiu for program support and formulating course design. This work was funded primarily by an NSF-NRT training grant (NSF 1828149) which established the Integrated training Model in Plant And Compu-Tational Sciences (IMPACTS) program at Michigan State University. This work is also supported by NSF Plant Genome Research Program awards IOS-2310355, IOS-2310356, and IOS-2310357. This project was supported by the USDA National Institute of Food and Agriculture, and by Michigan State University AgBioResearch and the Michigan State University College of Agriculture and Natural Resources Global Scholars program.

